# Mindfulness Training Alters Resting-State EEG Dynamics in Novice Practitioners via Mindful Breathing and Body-scan

**DOI:** 10.1101/2021.04.16.439387

**Authors:** Hei-Yin Hydra Ng, Changwei W. Wu, Feng-Ying Huang, Yu-Ting Cheng, Shiao-Fei Guu, Chih-Mao Huang, Chia-Fen Hsu, Yi-Ping Chao, Tzyy-Ping Jung, Chun-Hsiang Chuang

**Author notes:** Correspondence: Chun-Hsiang Chuang, PhD: No. 521, Nanda Road, Hsinchu, Taiwan 300193., and, Yi-Ping Chao, PhD: No. 259, Wenhua 1^st^ Rd., Guishan Dist., Taoyuan City, Taiwan, 333323. **Data availability** Following the regulations of Taiwan authority and the institutional review board, the data collected for this study are available only for the intended research topic by the corresponding personnel. **Conflict of interest disclosure** Declarations of interest conflict: none. **Ethics approval statement** This study was reviewed and approved by the Taipei Medical University Joint Institute Review Board (TMU-JIRB, project number N201905049). **Patient consent statement** All of the participants provided written informed consent prior to participation.

## Abstract

Mindfulness-based stress reduction (MBSR) has been proven to improve mental health and quality of life. This study examined how mindfulness training and various types of mindfulness practices altered brain activity. Specifically, the spectral powers of scalp electroencephalography (EEG) of the MBSR group who underwent an 8-week mindfulness training—including mindful breathing and body-scan—were evaluated and compared with those of the waitlist controls. Empirical results indicated that the long-term mindfulness intervention effect significantly elevated the resting-state beta powers and reduced resting-state delta powers in both practices; such changes were not observed in the waitlist control. Compared with mindful breathing, body-scanning resulted in an overall decline in EEG spectral powers at both delta and gamma bands among trained participants. Together with our preliminary data of expert mediators, the aforementioned spectral changes were salient after intervention, but mitigated along with expertise. Additionally, after receiving training, the MBSR group’s mindfulness and emotion regulation levels improved significantly, which were correlated with the EEG spectral changes in the theta, alpha, and low-beta bands. This study elaborated the neurophysiological correlates of mindfulness practices, suggesting that MBSR might function as a unique internal processing that involves increased vigilant capability and induces alterations similar to other cognitive training.

## 1. Introduction

Mindfulness refers to the mental state of being fully open and having attentional and nonjudgmental awareness of one’s internal and external experiences in the present moment (Kabat-Zinn, 1994). At present, mindfulness meditation has attracted global attention because of its benefits to practitioners’ mental health (Brown & Ryan, 2003). Mindfulness practices have been discovered to induce brain structure alterations (Fox et al., 2014), associated with improved working memory and attention (Mrazek, Franklin, Phillips, Baird, & Schooler, 2013; Van den Hurk, Giommi, Gielen, Speckens, & Barendregt, 2010). Mindfulness meditation can improve a practitioner’s self-regulation capability by increasing positive affect, life satisfaction, and well-being (Brown & Ryan, 2003; Garland, Geschwind, Peeters, & Wichers, 2015) and reducing depression, anxiety (Brown & Ryan, 2003; Davidson et al., 2003), stress (Irving, Dobkin, & Park, 2009), and even insomnia (Goldstein et al., 2019). Among various types of mindfulness interventions, mindfulness-based stress reduction (MBSR) is a standardized and secularized training program designed to improve mindfulness and coping abilities (Kabat-Zinn, 1994). MBSR programs typically span 8 weeks of weekly training, include one full-day workshop (Kabat-Zinn, 1994), and involve continuous mindfulness practices, such as mindful breathing, body-scan, and sitting meditation. Studies have demonstrated that after an 8-week training period, MBSR is generally effective in reducing depression and anxiety and promoting mental health (Fjorback, Arendt, Ørnbøl, Fink, & Walach, 2011). Although abundant evidence supports the role of MBSR in improving subjective perceptions, the brain mechanisms underlying MBSR remain to be investigated.

Mindfulness-based neuroscience studies have generally adopted a longitudinal approach instead of targeting situational practice effects. Objective measures of brain functions, such as electroencephalography (EEG) and functional magnetic resonance imaging (fMRI), have generally been adopted to test the efficacy of mindfulness interventions. For example, in a study of the EEG power of the experienced Rinpoche, with meditation experience of >10,000 hours, gamma-power enhancement was evident even during a resting state (Lutz, Greischar, Rawlings, Ricard, & Davidson, 2004), and this effect was sustained even during non-rapid-eye-movement (NREM) sleep (Ferrarelli et al., 2013). For meditation novices, EEG measures following an 8-week MBSR program have been widely associated with convergent and consistent outcomes. Researchers have found MBSR practitioners tend to exhibit stronger beta power in the frontal lobe during mindfulness practice than during the resting state (Gao et al., 2016). Similarly, MBSR practitioners exhibited elevated alpha power in the occipital and right temporal lobes (Ahani et al., 2014). Theta band power was reported to increase in the central, parietal, occipital, and left and right temporal lobes after the MBSR intervention by Ahani et al. (2014). Furthermore, MBSR practitioners exhibited lower delta power in the central–parietal area after MBSR intervention (Gao et al., 2016), and patients with chronic insomnia were also found to have lower delta power in the central lobe during NREM sleep after MBSR intervention (Goldstein et al., 2019). Overall, MBSR intervention is generally believed to enhance high-frequency EEG power (i.e., beta and gamma); however, its effect on low-frequency EEG power (i.e., theta and delta) remains uncertain.

Such neurophysiological evidence concerning MBSR is consistent with the evidence on stress reduction and cognitive improvement (Davidson et al., 2003). For example, the beta power in the frontal and temporal lobes of participants without stress stimuli was higher than that of the participants with stress stimuli (Hayashi et al., 2009), suggesting a negative relationship between beta power and stress level. As for the low-frequency bands, healthy adult participants with a high perceived stress level had higher delta and theta activity in the frontal, central, and parietal lobes, compared with those who had a low perceived stress level (Luijcks, Vossen, Hermens, van Os, & Lousberg, 2015). Another study on stress revealed that participants exhibited lower theta power under acute stressful conditions (Gärtner, Rohde-Liebenau, Grimm, & Bajbouj, 2014). Furthermore, another study highlighted how cognitive tasks elevated gamma power in comparison with the control conditions (Fitzgibbon, Pope, Mackenzie, Clark, & Willoughby, 2004), and a study on vigilance suggested that highly vigilant states corresponded to delta-power suppression (Smallwood & Schooler, 2015). Overall, EEG evidence generally reveals that stress reduction is positively correlated to low-frequency-band power and negatively correlated to high-frequency-band power, whereas cognitive performance and vigilance state are positively correlated to high-frequency-band power and negatively correlated to low-frequency-band power. EEG spectral powers can serve as objective functional markers of cognitive enhancement and stress reduction.

In addition to the long-term training effect of an 8-week MBSR program, the situational practice effect of such programs has recently attracted the attention of mindfulness researchers seeking to identify the variations between distinct mindfulness practices. The MBSR program involves a series of mindfulness practices (Davidson et al., 2003; Kabat-Zinn, 1994), such as mindful breathing, body-scan, compassion meditation, and open-monitoring. Davidson described the distinct practices associated with various cognitive effects in his book (Goleman & Davidson, 2017). Among the practices, mindful breathing and body-scan were most frequently used in previous studies (Ahani et al., 2014; Isbel, Lagopoulos, Hermens, & Summers, 2019; Wahbeh, Lane, Goodrich, Miller, & Oken, 2014), and both of them are associated with interceptive perceptions. However, a focus on breathing is a general practice in a variety of meditation approaches, whereas body-scan is specific to the MBSR program and is used to alleviate the chronic pain of clinical patients (Kabat-Zinn, 1990). The distinction between mindful breathing and body-scan can lead to different interoceptive effects on behavior and brain mechanisms. The fixed attention and relaxation in mindful breathing may differ from the attentional shifts to and from various body parts during body-scan. Recent studies have assessed the diverse effects of mindfulness practices using questionnaires and behavioral measures. One study demonstrated that people who practiced breath-focused meditation had a more nonjudgmental attitude toward themselves, showed more self-compassion, and experienced less emotional regulation difficulty, whereas those who practiced body-scan showed increased capabilities to describe their feelings and reduced rumination tendencies (Sauer-Zavala, Walsh, Eisenlohr-Moul, & Lykins, 2013). This suggests that mindful breathing and body-scan affect different brain functions. Some studies have suggested that body-scan yields more positive outcomes for practitioners than does breathing. For example, body-scan practice leads to a major increase in body awareness and a decrease in thought contents, whereas breathing practice engenders a comparatively less intense change (Kok & Singer, 2017). Another study on veterans with post-traumatic stress disorder disclosed that participants who practiced body-scan exhibited greater mindfulness improvement than did their breathing group counterparts (Colgan, Christopher, Michael, & Wahbeh, 2016). Studies with self-rating scales have established that body-scan seems to provide more promising benefits than mindful breathing. Nevertheless, the brain mechanisms targeted by the two mindfulness practices remain elusive at the current stage of research. Therefore, we adopted EEG measurement to examine the functional distinctions between mindful breathing and body-scan practices.

### 1.1 Working hypothesis

Collectively, the EEG spectral alterations associated with various mindfulness practices remain elusive, and whether the mindful breathing and body-scan practices take effect are yet to be tested. We first proposed that the functional distinction between these two mindfulness practices lies in the spatio-spectral disparity of the EEG, meaning that frontal power elevation follows body-scan, and parietal power reduction follows mindful breathing after the 8-week MBSR intervention. Second, we proposed that the functional distinction between practices is amplified by the long-term training effect (not observed in novices). Accordingly, we designed an EEG experiment to assess the neurophysiological changes in terms of both situational mindfulness practice and long-term training effects. In addition, we conducted the same protocol with a waitlist control group without MBSR training to enable cross-group comparison.

## 2. Materials and Methods

### 2.1 Participants

Forty-three volunteers were invited to participate in the MBSR training and register for an 8-week MBSR course in the fall of 2019. Before the course started, all volunteers were required to undergo one of the three orientation sessions to understand the details of the procedure, compensation, potential risks, and the contributions of this study. Potential participants were screened by applying the following exclusion criteria: being outside the age range of 20 to 80 years old; having prior experience of mindfulness meditation; having a metabolic illness, any history of mental illness, neural illness, or epilepsy; being a smoker or drug addict; and having any bodily metallic implant, claustrophobia, or pregnancy. Ten people who did not meet the criteria were excluded, and eventually 33 (4 men and 29 women) people aged between 29 and 68 years participated in the study (mean = 47.46, SD = 8.79; 18 participants in the MBSR group, 15 participants in the waitlist control group). The participants were new to mindfulness training before the study, and they received a free 8-week MBSR course for electing to participate. All of the participants provided written informed consent prior to participation. This study was reviewed and approved by the Taipei Medical University Joint Institute Review Board (TMU-JIRB, project number N201905049). Following the regulations of Taiwan authority and the institute review board, the data collected for this study are available only for the intended research topic by the corresponding personnel.

### 2.2 Intervention: MBSR and waitlist control

Each participant completed the EEG–fMRI experiments twice. For the MBSR group, two experiments were scheduled before and after the 8-week MBSR training (pre-test and post-test). For the waitlist control group, two scans were performed 8 weeks apart (same pre-test and post-test); however, they were prohibited from accessing and receiving information regarding mindfulness practice during this period. The MBSR group participated in an 8-week standardized MBSR course, as proposed by Santorelli, Kabat-Zinn, Blacker, Meleo-Meyer, and Koerbel (2017). The MBSR course was instructed by a licensed MBSR instructor. MBSR classes were conducted weekly for 2.5 h for 8 weeks in addition to a one-day mindfulness workshop. These weekly meetings involved the development of various mindfulness skills, dialogue and reflection on mindfulness home practice, and practice segments. The participants performed types of mindfulness practice such as a sitting meditation entailing breathing, mindful listening, body-scan, and mindful yoga. Participants were also assigned daily homework comprising both formal and informal meditation activities. Formal activities, including body-scan practice, sitting meditation, mindful yoga, mountain/lake meditation, or loving kindness meditation, required 45 min to complete each day. The participants were asked to complete a practice and information sheet. The participants in the waitlist control group were instructed to maintain their usual life activities but not engage with any mindfulness-related information during the 8 weeks. After the posttest experiment, the waitlist control group participants underwent an MBSR program for compensation.

### 2.3 Experimental procedure

The study procedure is presented in Fig. 1. In the orientation sessions, the study was explained to the participants; in particular, the use of the fMRI machine was detailed, as this study employed the EEG–fMRI simultaneous scanning technique (however, only EEG and behavioral information are included in this report). Thereafter, the participants finished the informed consent form and a questionnaire battery. The questionnaires were the Five Facet Mindfulness Questionnaire (FFMQ)(Baer et al., 2008)—Taiwanese version, the Difficulties in Emotion Regulation Scale (DERS)(Gratz & Roemer, 2004), and demographic questionnaires. The questionnaire battery took approximately 30 minutes to complete.

**Figure 1.**
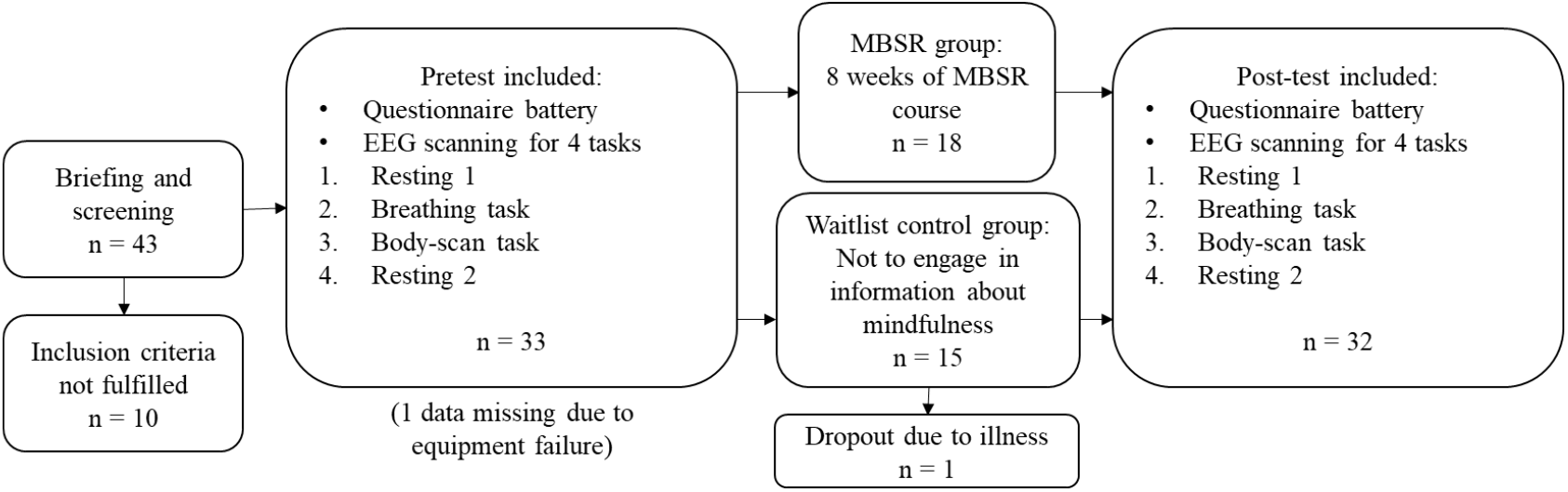
Study design and recruitment procedure

After the orientation sessions and within 2 months before the MBSR course, the researchers made an appointment for the participants to come into the laboratory for the pretest experiment. The participants were introduced to the tasks that they would perform during the scanning while an EEG cap was being set up. In the scanning section, participants laid down in the MRI machine to perform four tasks. A mirror setting was installed with projectors for visual presentations. The first task was the resting state (***resting 1***): Participants were instructed not to think of anything specific with their eyes closed, not to move their heads, and not to fall asleep. The second task was the mindful breathing (***breathing***): Participants were instructed to pay attention to their natural breath and focus on the sensation near their noses during respiration. Whenever they realized that they were getting distracted, they were instructed to press a button on the right hand. The third task was the body-scan (***body-scan***): Participants were instructed to perceive the most salient body sensations and shift focus between various body parts during the session. Similar to the breathing task, they were instructed to press a button when they were distracted. Finally, an additional resting state (***resting 2***), identical to ***resting 1***, was conducted to evaluate whether the brain status was returned to baseline after the mindful practices. Every task lasted 5 minutes, and if participants opened their eyes during a task, they would see a white fixation cross on a black background. We used E-Prime 2.0.10 (Psychology Software Tools, Pittsburgh, PA, USA) for instruction presentation and response recording. After the pre-test experiment, the MBSR group participated in an 8-week MBSR course while the waitlist control group went about their usual life. Within 3 months after the completion of the MBSR course, all participants from both groups participated in the post-test experiments that were identical to the pre-test sessions.

### 2.4 Questionnaires

#### 2.4.1 The Five Facet Mindfulness Questionnaire (FFMQ)

is a self-reported assessment for measuring mindfulness level (Baer et al., 2008). The FFMQ concerns five aspects of mindfulness, namely observing, describing, being self-aware, having a nonjudgmental attitude toward inner experience, and being nonreactive to inner experience. Sample questions of the FFMQ are “I notice the smells and aromas of things” (observing), “I am good at finding words to describe my feelings” (describing), “I find myself doing things without paying attention (reversed)” (self-awareness), “I think some of my emotions are bad or inappropriate and I should not feel them (reversed)” (nonjudgmental), and “I perceive my feelings and emotions without having to react to them” (nonreactive). The FFMQ has 39 items, of which 19 are reversed. Items are rated on a 5-point Likert scale, ranging from 1 (*never or very rarely true*) to 5 (*very often or always true*), and a total score from 39 to 195 can be obtained after answering the whole questionnaire. The FFMQ has good reliability (α= .72 to .92; Baer et al., 2008). In this study, we employed a Taiwanese version of the FFMQ that was in Chinese (F. Y. Huang et al., 2015), and its reliability was also satisfactory (α = .91 and α = .96 in the pre-test and post-test, respectively).

#### 2.4.2 The Difficulties in Emotion Regulation Scale (DERS)

is a self-reported measurement tool for examining the level of difficulty experienced by people in addressing their negative emotions and producing desirable outcomes (Gratz & Roemer, 2004). The DERS concerns six aspects of emotion regulation difficulties, namely nonacceptance of emotional responses, difficulties engaging in goal-directed behaviors, impulse control difficulties, lack of emotional awareness, limited access to emotion regulation strategies, and lack of emotional clarity. Sample items are “When I’m upset, I feel guilty for feeling that way” (nonacceptance), “When I’m upset, I have difficulty getting work done” (goals), “When I’m upset, I lose control over my behaviors” (impulse), “I pay attention to how I feel (reversed)” (awareness), “When I’m upset, I believe that I will remain that way for a long time” (strategies), and “I am confused about how I feel” (clarity). The DERS has 36 items, of which 11 are reversed. Items are rated on a 5-point Likert scale, ranging from 1 (*almost never*) to 5 (*almost always*), and the total score obtained ranges from 36 to 180. The DERS has good reliability (α = .80 to .89). This study employed a Taiwanese version of the DERS that was translated into Chinese through the back-translation procedure, and the reliability analysis yielded α = .96 and α = .95 for the pre-test and post-test, respectively.

#### 2.4.3 EEG measurement and analysis

In each experiment, simultaneous EEG–fMRI signals were recorded for each functional scan using a 3T PRISMA MRI scanner (Siemens, Erlangen, Germany). The 32-channel EEG data were recorded using an MR-compatible system (Brain Products GmbH, Gilching, Germany) that was positioned according to the international 10/20 system. The built-in impedance in each electrode was 5 kΩ, and abrasive electrode paste (Abralyt HiCl) was used to reduce the electrode–skin impedance. The EEG signal was recorded synchronously with the MR trigger using Brain Vision Recorder (Brain Products) with a 5k-Hz sampling rate and a 0.5 μV voltage resolution (reference at FCz). A band-pass filter was set with cut-off frequencies of 250 Hz and 0.0159 Hz, and an additional 60-Hz notch filter was employed. Here, we have reported only the EEG outcomes, as the MRI results were designed to be reported separately.

Recorded EEG data were re-sampled to 50,000 Hz and then corrected for gradient-induced artifacts. Ballistocardiographic artifacts were corrected using the adaptive average subtraction method, and the R-peak intervals were estimated from the electrocardiogram electrode through Analyzer 2.1 (Brain Products) after the data were down-sampled to 250 Hz. The EEG data were then filtered with a 0.2–40 Hz band-pass FIR filter. Thereafter, an independent component analysis (ICA, with Infomax method) to eliminate artifacts caused by electrooculogram (EOG), electromyogram (EMG), and electrocardiogram (ECG) artifacts and the remaining MRI-induced artifacts. The artifact-free data were referenced to an average electrode across whole scalps, as recommended by J. J. B. Allen, Coan, and Nazarian (2004) and Davidson (1998). The processed data were converted into frequency domain representations using short-time Fourier transformation and Welch’s periodogram method. Specifically, each 5-minute EEG signal (75,000 points) was divided into 256-point segments using the Hanning window with 128 points overlapped. Each segment was zero-padded to 512 points, followed by a 512-point fast Fourier transformation. All the resultant spectra were subsequently log-transformed (10log10, results in dB; J. J. B. Allen et al., 2004) and averaged over segments. Finally, average EEG band powers were calculated in the frequency ranges of 1–4 Hz, 4–8 Hz, 8–13 Hz, 13–20 Hz, 20–30 Hz, and 30–40 Hz, representing delta, theta, alpha, low-beta, high-beta, and gamma band powers, respectively (Abhang, Gawali, & Mehrotra, 2016; Deuschl & Eisen, 1999; Hauswald, Übelacker, Leske, & Weisz, 2015; Rangaswamy et al., 2002; Teplan, 2002). These signal processes were conducted using the EEGLAB toolbox 13.6.5b (Swartz Center for Computational Neuroscience, University of California San Diego; Delorme & Makeig, 2004) with Matlab R2019a.

### 2.5 Statistical analysis

To ensure that all parameters (EEG band power and FFMQ and DERS scores) were fitted for parametric analyses, Kolmogorov–Smirnov tests were performed to examine normality. The chi-square test and *t* test were conducted to examine whether demographic features, FFMQ and DERS levels were different between the MBSR group and the waitlist control group before the MBSR intervention. Subsequently, two more *t* tests were performed to examine whether a significant difference existed in terms of FFMQ and DERS in both groups before and after the MBSR intervention.

To examine whether EEG power spectra changed before and after the MBSR intervention, a paired-sample *t* test was performed for every 0.5-Hz frequency bin in all four tasks (***resting 1, breathing, body-scan, resting 2***) in both groups and for both channels, Fz and Pz. Furthermore, Pearson’s correlation analysis was performed to examine whether a correlation existed between the change of EEG band power before and after the MBSR intervention and the change of FFMQ and DERS scores in both groups. The change of EEG power was calculated by subtracting the pretest EEG band power from the posttest EEG band power (post-test – pre-test), and the change in FFMQ and DERS scores was calculated similarly.

## 3. Results

### 3.1 Normality test

Table 1 presents the mean FFMQ and DERS scores rated by both the MBSR and control groups during the pretest and post-test sessions. The MBSR group (N = 18) scored 114.94 ± 16.99 on the FFMQ_pre_, 142.56 ± 23.24 on the FFMQ_post_, 97.78 ± 20.72 on the DERS_pre_, and 84.72 ± 23.21 on the DERS_post_. The control group (N = 15) scored 116.80 ± 16.45 on the FFMQ_pre_, 115.87 ± 14.65 on the FFMQ_post_, 84.87 ± 22.53 on the DERS_pre_, and 98.80 ± 16.12 on the DERS_post_. All these scores validated the assumption of normality (Kolmogorov–Smirnov test: *ps* > .05). In terms of resting-state EEG activity, the logarithmic powers of the delta, theta, alpha, low-beta, high-beta, and gamma bands over channels Fz and Pz recorded in the two sessions for both groups also validated the assumption of normality (Kolmogorov–Smirnov test: *ps* > .05). The only exception was the posttest gamma score for channel Pz in the waitlist control group (*p* = .032). Because our analysis focused on the main effects of the MBSR intervention group, parametric tests were employed in the subsequent analyses.

**Table 1.** Demographic data and questionnaire statistics

|  | Waitlist control group | MBSR group | Statistics | *p* |
| --- | --- | --- | --- | --- |
| N | 15 | 18 |  |  |
| Gender (female) | 13 | 17 | $\chi^2(1) = 1.60$ | 0.206 |
| Age | 46.67±8.03 | 48.50±9.52 | $t(31) = 0.59$ | 0.559 |
| Year of Education | 17.60±2.53 | 16.89±2.03 | $t(31) = -0.90$ | 0.377 |
| FFMQ<sub>pre</sub> | 116.80±16.45 | 114.94±16.99 | $t(31) = -0.32$ | 0.753 |
| FFMQ<sub>post</sub> | 115.87±14.65 | 142.56±23.24 | $t(17) = 5.32$ | <.001 |
|  | -0.93±13.62 | 27.61±22.02 |  |  |
| FFMQ<sub>post-pre</sub> | $t(14) = -0.27$<br>$p = 0.796$ | $t(17) = 5.26$<br>$p < .001$ | | |
| DERS<sub>pre</sub> | 94.87±22.53 | 97.78±20.72 | $t(31) = 0.39$ | 0.702 |
| DERS<sub>post</sub> | 98.80±16.12 | 84.72±23.21 | $t(31) = -2.08$ | 0.046 |
|  | 3.93±19.97 | -13.06±25.87 |  |  |
| DERS<sub>post-pre</sub> | $t(14) = 0.76$<br>$p = 0.458$ | $t(17) = -2.14$<br>$p = 0.047$ | | |

*Note: FFMQ: Five Facet Mindfulness Questionnaire; DERS: Difficulties in Emotion Regulation Scale*

### 3.2 Demographic and behavioral analysis

Thirty-three participants (29 women and 4 men, aged 47.67 ± 8.79 years) were recruited in this study, of whom 15 (12 women and 3 men, aged 46.67 ± 8.03 years) served as the control group and 18 (17 women and 1 man, aged 48.50 ± 9.52 years) served as the MBSR group. As evident in the demographic data and questionnaire results presented in Table 1, no significant difference was apparent in terms of gender (χ^2^ [1] = 1.60, *p* = .206), age (*t* [31] = 0.59, *p* = .559), and educational level (*t* [31] = −0.90, *p* = .377) between the waitlist control and MBSR groups.

A between-group analysis showed that the differences between the two groups in both questionnaire responses were significant in the post-test session (FFMQ: *t* [31] = 5.32, *p* < .001; DERS: *t* [31] = −2.08, *p* = .046) but not in the pretest session (FFMQ: *t* [31] = −0.32, *p* = .753; DERS: *t* [31] = 0.39, *p* = .702). A within-group analysis showed that after the MBSR intervention, the MBSR group exhibited a significant increase in the FFMQ score (*t* [17] = 5.32, *p* < .001) and a decrease in the DERS score (*t* [17] = −2.14, *p* = .047) from the scores before the intervention, whereas the FFMQ (*t* [14] = −0.27, *p* = .796) and DERS (*t* [14] = 0.76, *p* = .458) scores of the waitlist control group remained unchanged.

### 3.3 EEG comparison between pretest and post-test sessions

After an 8-week mindfulness training, the effect of mindfulness practices on resting-state EEG activity was examined. Figures 2 and 3 illustrate the comparisons of power spectrum density (PSD) between the pretest and post-test for Fz and Pz electrodes, respectively; the black asterisks indicate the significant differences in EEG powers identified at certain frequency bins. Student’s *t* test was applied for every 0.5-Hz frequency bin ranging from 0 Hz to 40 Hz (0–0.2 Hz was filtered out during preprocessing), followed by false discovery rate (FDR) correction for conducting multiple comparisons across frequency bins and channels.

**Figure 2.**
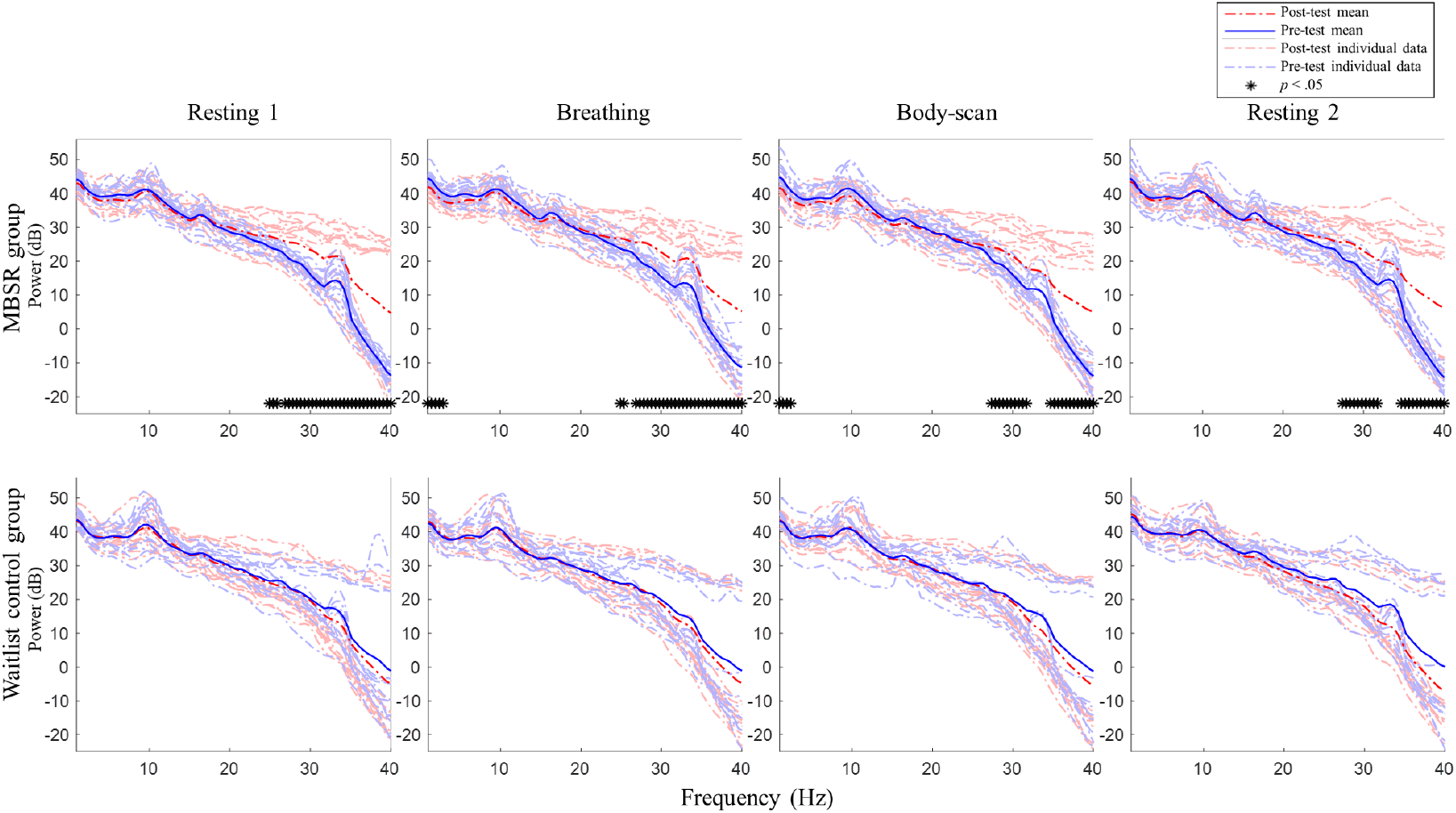
Post-pretest PSD t-test for channel Fz between the MBSR group and waitlist control. MBSR group n = 17, waitlist control group n = 14. The statistics were corrected for false discovery rate (FDR).

**Figure 3.**
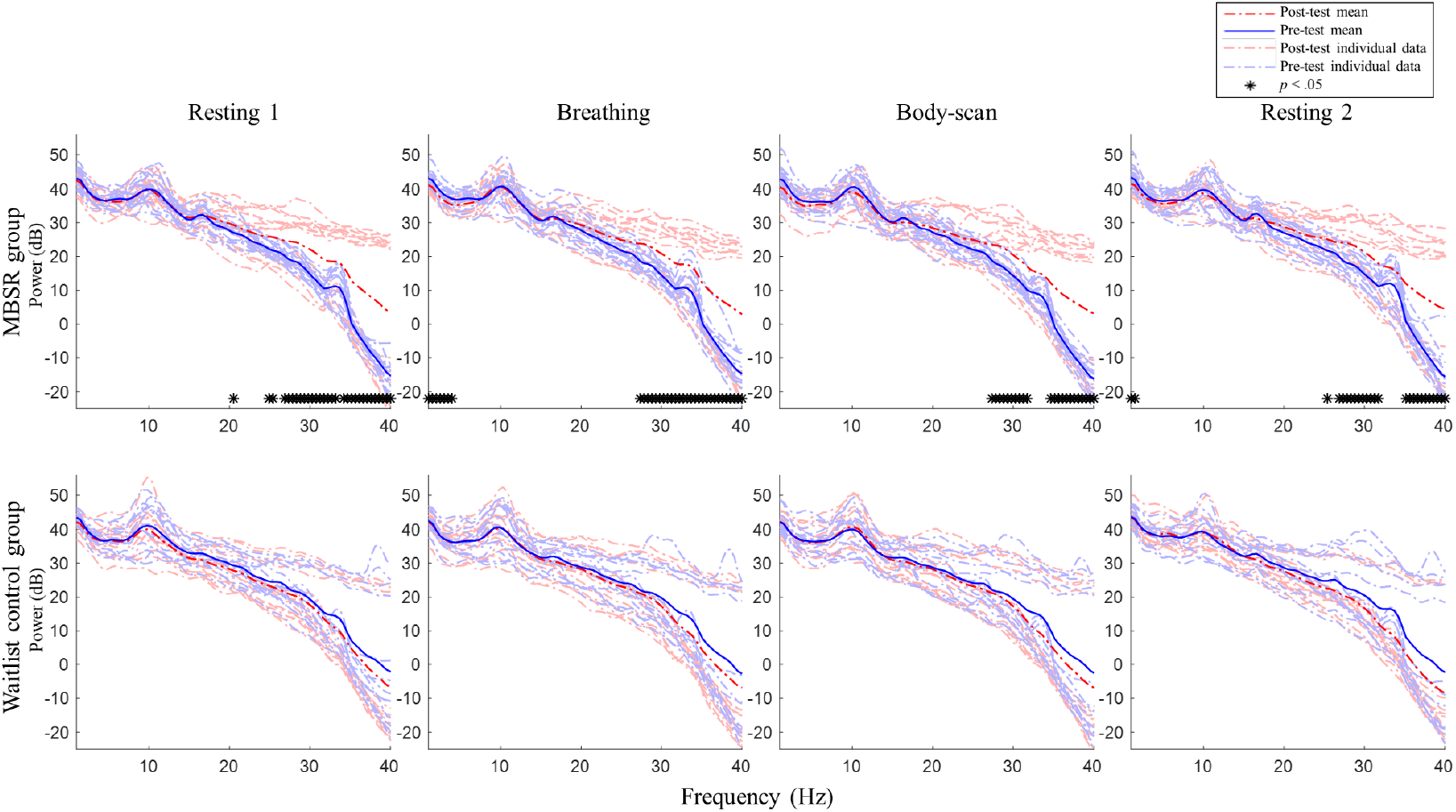
Post-pretest PSD t-test for channel Pz between the MBSR group and waitlist control. MBSR group n = 17, waitlist control group n = 14. The statistics were corrected for false discovery rate (FDR).

For the MBSR group, compared with the pretest session, the EEG activity of Fz and Pz sites measured in the posttest session revealed the following: 1) in ***resting 1***, high-beta and gamma powers significantly increased (Fz: 24.5–40 Hz except 26 Hz, *ts* ≥ 2.56, *ps* < .05; Pz: 20–20.5 Hz and 24.5–40 Hz, except 25.5, 26, and 35.5 Hz, *ts* ≥ 2.41, *ps* < .05); 2) in ***breathing***, high-beta and gamma powers significantly increased (Fz: 24.5–40 Hz except 25.5–26 Hz, *ts* ≥ 2.31, *ps* < .05; Pz: 27–40 Hz, *ts* ≥ 2.70, *ps* < .05) and delta powers significantly decreased (Fz: 0.2–2 Hz, *ts* ≥ −2.72, *ps* < .05; Pz: 0.2–3 Hz, *ts* ≥ −2.61, *ps* < .05); 3) in ***body-scan***, high-beta and gamma powers significantly increased (Fz: 27–31.5 Hz, *ts* ≥ 2.83, *ps* < .05; 34.5–40 Hz, *ts* ≥ 2.74, *ps* < .05; Pz: 27–31.5 Hz, *ts* ≥ 3.05, *ps* < .05; 34.5–40 Hz, *ts* ≥ 2.94, *ps* < .05) and delta power significantly decreased only at Fz (0.2–1.5 Hz, *ts* ≥ −2.76, *ps* < .05); 4) in ***resting 2***, high-beta and gamma powers significantly increased (Fz: 27–31.5 Hz, *ts* ≥ 2.80, *ps* < .05; 34.5–40 Hz, *ts* ≥ 2.85, *ps* < .05; Pz: 25–31.5 Hz except 25.5 and 26 Hz, *ts* ≥ 2.99, *ps* < .05; 35–40 Hz, *ts* ≥ 3.56, *ps* < .05) and delta powers significantly decreased only at Pz (0.2–0.5 Hz, *ts* ≥ −2.80, *ps* < .05). For the waitlist control group, no significant difference was observed in EEG power between the posttest and pretest sessions in all four tasks (lower panel of Figs. 2 and 3, *ps* > .05).

Figure 4 presents the spatial distribution of the EEG spectral power differences between the post-test and pre-test of the MBSR group in the four tasks. Paired-sample *t* tests were applied to all channels to compare the difference between the post-test and -test conditions. The channel locations that do not exhibit significant differences (*ps* > .05) were marked as zero difference and depicted in green in the topoplots. The results showed that the high-beta and gamma bands had significant spectral differences across the whole scalp in ***resting 1*** and ***resting 2***. There were also some small-scale differences in the delta and low-beta bands in the lateral frontal, left parietal, and right temporal areas in ***resting 1*** and ***resting 2***. The delta band had significant EEG power differences in the frontal and parietal areas, and the high-beta and gamma bands had significant power differences across the whole scalp in the ***breathing*** and ***body-scan*** tasks. There were also significant differences in the theta band in the frontal and occipital areas in the ***breathing*** and ***body-scan*** tasks. For the ***breathing*** task, there are small-scale differences in the alpha and low-beta bands in the right temporal area. For the ***body-scan*** task, there a small-scale difference in the alpha in the right temporal area, and a frontal power asymmetry (right > left) in the low-beta band.

**Figure 4.**
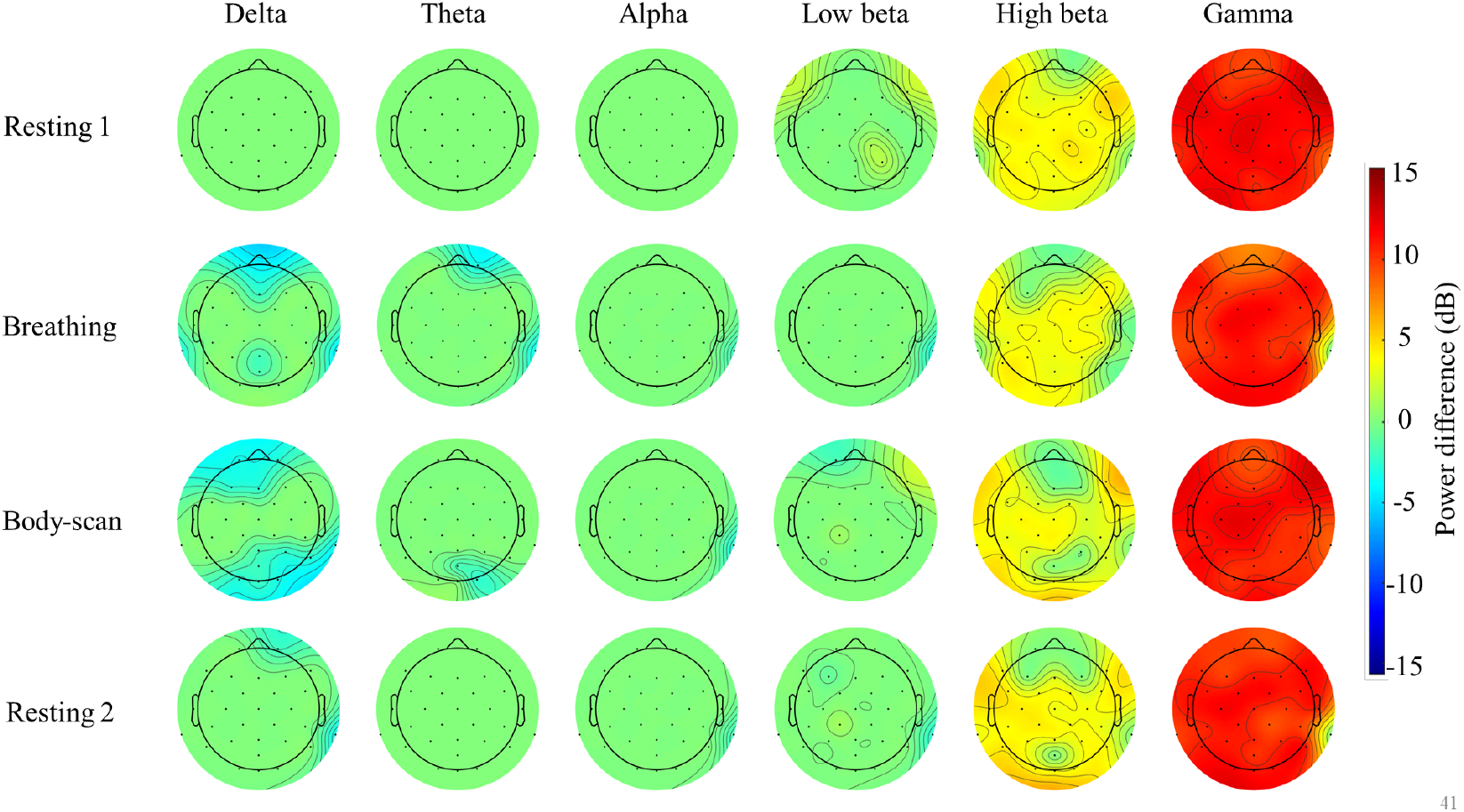
Topoplot analysis of the EEG spectral power of the MBSR group in the breathing and body-scan tasks, and the paired t-test comparison between post-test and pretest. Power difference = Post-test – pre-test power. Channel locations with zero power difference represent no significant difference (paired-sample *t* test, *ps* > .05, depicted in green)

### 3.4 EEG correlates of mindfulness practice

This study further investigated the momentary state effect of acquired mindfulness skills by examining the EEG activity. To this end, the EEG powers during ***breathing*** and ***body-scan*** were referenced to that during ***resting 1***. Figure 5 shows the spectral comparisons of ***breathing, body-scan***, and ***resting 2*** in the post-test session. Compared with ***resting 1***, the powers of delta, low-beta, high-beta, and gamma bands significantly decreased during ***body-scan*** at both Fz (0.2–2.5 Hz, *ts* ≥ −2.55, *ps* < .05; 15–17.5 Hz, *ts* ≥ −2.92, *ps* < .05; 19.5–20.5 Hz, *ts* ≥ −2.59, *ps* < .05; 23.5– 24.5 Hz, *ts* ≥ −2.54, *ps* < .05; 28.5–37 Hz, *ts* ≥ −2.59, *ps* < .05) and Pz (0.2–4 Hz, *ts* ≥ −3.08, *ps* < .05; 13–16.5 Hz, *ts* ≥ −3.00, *ps* < .05; 29.5–37.5 Hz, *ts* ≥ −2.44, *ps* < .05). However, no significant power change was found in ***breathing*** and ***resting 2*** (*ps* > .05). The EEG power during ***body-scan*** was also referenced to the power during ***breathing***, and no significant difference was found (*ps* > .05). For the waitlist control group that did not engage in any mindfulness practice, no significant differences were found in the EEG power between every task pair (*ps* > .05).

**Figure 5.**
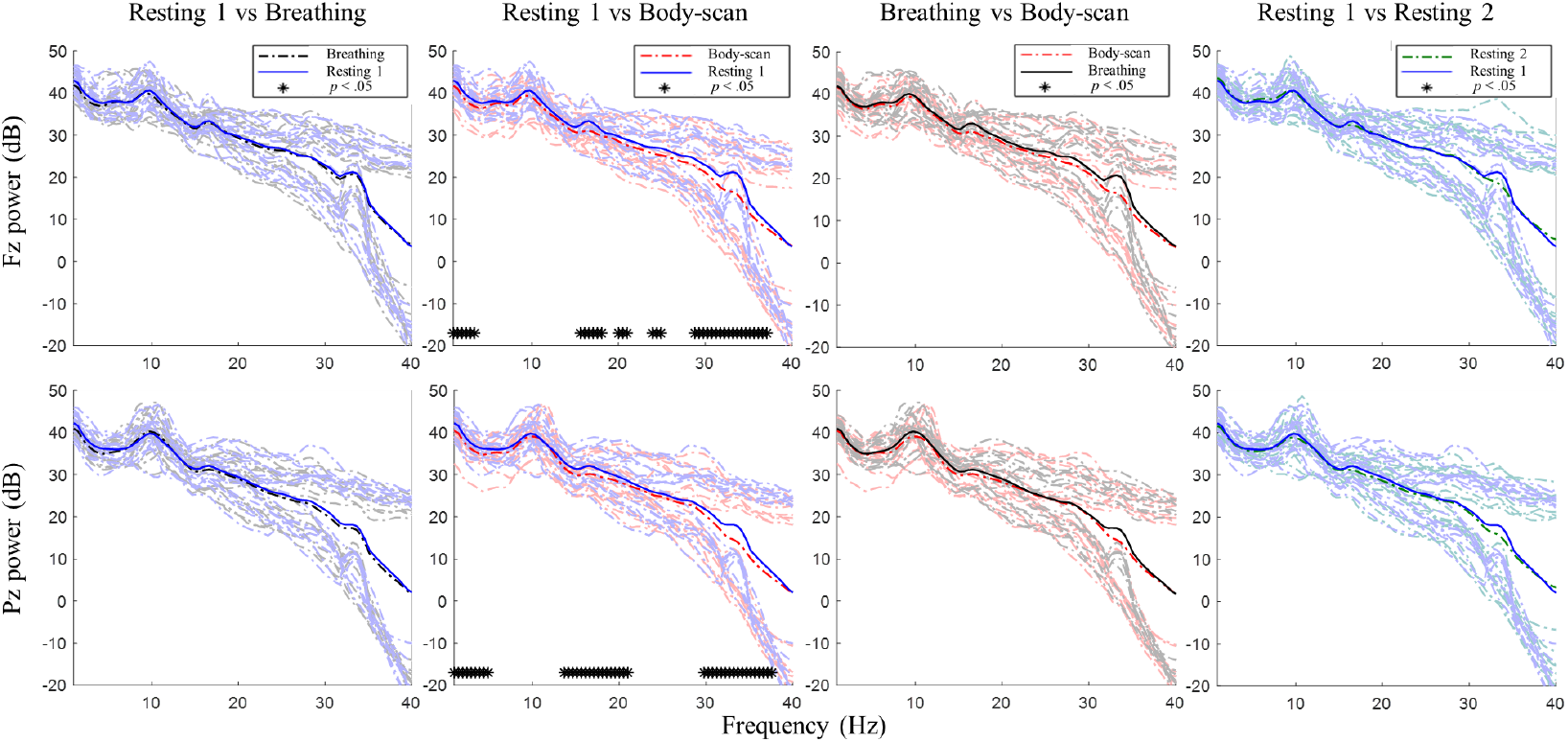
Post-test PSD t-test among the four tasks in the MBSR group. n = 18. No significant difference was found in the posttest for the waitlist control group. No significant difference was found in the pretest for either group.

### 3.5 EEG correlates of mindfulness and behavioral measures

Correlation analysis was performed to examine the potential correlations between EEG activities and behavioral outcomes. Table 2 presents the correlation coefficients between the post-test–pre-test differences of EEG power in the four tasks and those in the behavioral measures. For the MBSR group, Δlow-beta power in ***resting 1*** for the Pz site was found to have a negative correlation with ΔDERS (*b* = −0.50, *p* = .040), Δtheta in ***breathing*** for the Pz site had a marginally significant negative correlation with ΔFFMQ (*b* = −0.48, *p* = .053), Δlow beta in ***body-scan*** for the Fz site had a marginally significant negative correlation with ΔFFMQ (*b* = −0.48, *p* = .054), Δalpha in ***resting 2*** for both Fz (*b* = −0.64, *p* = .006) and Pz sites (*b* = −0.51, *p* = .036) showed a negative correlation with ΔFFMQ, and Δtheta in ***resting 2*** for the Pz site had a marginally significant negative correlation with ΔFFMQ (*b* = −0.47, *p* = .058).

**Table 2.** Correlation coefficients among the changes of EEG wave bands and behavioral measures in MBSR group

| Fz | | Δdelta | Δtheta | Δalpha | Δlow beta | Δhigh beta | Δgamma |
| --- | --- | --- | --- | --- | --- | --- | --- |
| Resting 1 | ΔFFMQ | -0.06 | 0.02 | -0.20 | 0.16 | 0.16 | 0.15 |
| | ΔDERS | 0.14 | 0.07 | 0.31 | -0.08 | -0.24 | -0.19 |
| Breathing | ΔFFMQ | -0.31 | -0.15 | -0.12 | -0.09 | 0.02 | 0.11 |
| | ΔDERS | 0.25 | 0.12 | 0.05 | 0.10 | -0.19 | -0.18 |
| Body-scan | ΔFFMQ | -0.01 | -0.28 | -0.36 | -0.48 † | 0.14 | 0.16 |
| | ΔDERS | -0.08 | 0.24 | 0.16 | 0.40 | -0.23 | -0.23 |
| Resting 2 | ΔFFMQ | -0.25 | -0.28 | -0.64** | -0.07 | 0.21 | 0.17 |
| | ΔDERS | 0.42 | 0.34 | 0.37 | -0.15 | -0.33 | -0.20 |
| Pz | | Δdelta | Δtheta | Δalpha | Δlow beta | Δhigh beta | Δgamma |
| Resting 1 | ΔFFMQ | -0.06 | 0.03 | -0.10 | 0.38 | 0.25 | 0.34 |
| | ΔDERS | -0.04 | -0.07 | -0.03 | -0.50* | -0.42 | -0.42 |
| Breathing | ΔFFMQ | -0.29 | -0.48 ‡ | -0.25 | -0.12 | 0.20 | 0.31 |
| | ΔDERS | 0.07 | 0.23 | -0.04 | -0.05 | -0.39 | -0.38 |
| Body-scan | ΔFFMQ | -0.09 | -0.35 | -0.41 | -0.03 | 0.36 | 0.30 |
| | ΔDERS | 0.06 | 0.25 | 0.30 | -0.05 | -0.44 | -0.42 |
| Resting 2 | ΔFFMQ | -0.20 | -0.47§ | -0.51* | -0.06 | 0.22 | 0.15 |
| | ΔDERS | 0.21 | 0.22 | 0.17 | -0.25 | -0.38 | -0.18 |

Notes: \* $p < 0.05$ \*\* $p < 0.01$ . $\Delta$ = post-test score – pre-test score. †, ‡, § = marginally significant, $p = .054 / .053 / .058$ respectively. For the waitlist control group, none of the post-pretest difference of EEG power was significantly correlated with that of any behavioral measure, neither for Fz nor Pz site.

## 4. Discussion

### 4.1 Summary of results

Table 1 presents the MBSR intervention effects on the FFMQ and DERS scores, indicating that participants demonstrated augmented levels of trait mindfulness and emotional regulation. The EEG spectral results (Figs. 2 and 3) showed that the 8-week intervention led to an increase in high-frequency EEG activities across all conditions and a decrease in low-frequency EEG activities during the ***breathing*** and ***body-scan*** practices. As observed in the MBSR group at the follow-up visit in week 8 (Fig. 5), ***body-scan*** suppressed EEG power across all frequencies. Additionally, the mindfulness-induced changes in theta, alpha, and low-beta band powers significantly correlated with the changes in FFMQ or DERS scores.

### 4.2 Training effect of MBSR

The differences between the post-test and pre-test results in ***resting 1*** indicated the long-term effect of the MBSR practice on EEG activity. Previous studies have reported that relaxation exercises induced decreased beta and gamma band power (Stinson & Arthur, 2013) and increased theta band power (Field, Diego, & Hernandez-Reif, 2010). The current study’s EEG results and previous research on MBSR (Cahn, Delorme, & Polich, 2010; Lutz et al., 2004) have demonstrated that, unlike relaxation exercises, mindfulness practices yield increased high-frequency EEG activity. This long-term neuro-electrical change was sustained in our study irrespective of which mindfulness task the participants engaged in. In the MBSR group, the post-pretest EEG power comparisons of ***resting 1, breathing, body-scan***, and ***resting 2*** revealed the same pattern, namely elevated high-frequency (high beta and gamma) EEG power in the post-test EEG scan. The results further supported our argument that the increase in high-frequency EEG power is a long-term effect of mindfulness practice. Although mindfulness practice is known to provide the same relaxation effect as other relaxation exercises (Davidson et al., 2003), this study suggested that mindfulness practice differs from relaxation exercises.

The results of mindfulness practice were rather similar to the EEG findings on beta and gamma neurofeedback training (a training aimed at improving beta or gamma power) that showed that 10 days of beta and gamma neurofeedback training led to improved episodic memory among healthy adults (Keizer, Verschoor, Verment, & Hommel, 2010). A similar association between long-term cognitive training and gamma power elevation was also found among the elderly population (Staufenbiel, Brouwer, Keizer, & Van Wouwe, 2014) and patients with Alzheimer’s disease (Van Deursen, Vuurman, Verhey, Van Kranen-Mastenbroek, & Riedel, 2008), schizophrenia (Molina et al., 2020), and attention-deficit/hyperactivity disorder (Yordanova, Banaschewski, Kolev, Woerner, & Rothenberger, 2001). Furthermore, the results were consistent with the neuroimaging findings that highly focused participants with 6 weeks of mindfulness training showed better cognitive performance in the Stroop test, in addition to exhibiting higher dlPFC activation (M. Allen et al., 2012). With results showing elevated high-frequency EEG power, mindfulness practice is likely similar in effect to cognitive training.

After the MBSR intervention, the low-frequency (delta) activity at Fz decreased during ***breathing*** and ***body-scan*** (Fig. 2) and that at Pz decreased during ***breathing*** (Fig. 3). Previous studies on different mindfulness styles (i.e., Vipassana and Qigong) have suggested that compared with non-meditators, practitioners’ delta activity increased in the prefrontal area during the resting state (Cahn et al., 2010; Tei et al., 2009). However, another study on MBSR reported that practitioners’ frontal delta power decreased (Gao et al., 2016), indicating that different mindfulness practices may lead to different EEG results. Our data supported the results of Gao et al. (2016) that MBSR practitioners’ low-frequency EEG power decreases and high-frequency EEG power increases. The current study further suggested that such EEG power changes could be introduced with 8 weeks of training.

Some have argued that the delta oscillation of EEG is related to mental task performance because delta oscillation represents people’s attention to their internal process (Harmony, 2013; Harmony et al., 1996). Knyazev (2007) further suggested that delta and alpha oscillations in the prefrontal area may contribute to a reciprocal inhibitory mechanism that can manipulate people’s motivation and attention and moderate people’s mental task performance. Harmony (2013) suggested that the increased delta oscillation in mindfulness practitioners represents the inhibition of the prefrontal cortex in addition to the reduction of emotional and cognitive engagement. MBSR practitioners, however, are instructed to not inhibit their emotions or cognition but to accept and merely observe their inner thoughts just as they are (Davidson et al., 2003). This practice of acceptance may explain the current result that delta power decreased during mindfulness practice after 8 weeks of training. This argument is further legitimized by the result that only the MBSR-trained participants, and not the waitlist control group, demonstrated the delta power drop. This is because attentional yet nonjudgmental acceptance of oneself requires continuous and effort-intensive practices to master.

Additionally, this delta power suppression suggested that the practices might involve a stage of high vigilance (Smallwood & Schooler, 2015). A previous study on mind-wandering (Braboszcz & Delorme, 2011) showed that when people were distracted from a task and started mind-wandering, their delta and theta power increased whereas their alpha and beta power dropped. The EEG results of mind-wandering were exactly the opposite of our mindfulness results of decreased delta power. The contradictory EEG results support the argument that mindfulness is a process of disciplining the mind and stopping mind-wandering (mindlessness; Davidson et al., 2003; Mrazek, Smallwood, & Schooler, 2012). Furthermore, another study suggested that mindfulness programs can reduce mind-wandering episodes (Schooler et al., 2014). This claim is supported by the results that participating in 8 weeks of mindfulness training promotes working memory capacity, which is the key to maintaining focus in a cognitively demanding and vigilance-requiring situation (Jha, Stanley, Kiyonaga, Wong, & Gelfand, 2010). Even 8 minutes of mindful breathing was found to reduce people’s attentional error in a vigilance task (Mrazek et al., 2012). Therefore, EEG spectral whitening (low-frequency power drop and high-frequency power elevation) represents a mental state with less mind-wandering and higher mindfulness.

Figure 4 plots the spatial distributions of the post-test – pre-test EEG spectral differences of the four tasks. Our results echoed the literature review that the changes in mindfulness-induced EEG power were evident in the frontal and parietal areas. Figure 4 further suggested that some other brain areas, including the occipital and temporal areas, also exhibit comparable spectral changes. Also, there was a frontal power asymmetry in the low beta-band in ***body-scan***. We expect that further studies on the frontal EEG power asymmetry will be legitimate in the exploration of mindfulness (also see Isbel et al., 2019).

It is noted that some participants in the MBSR group did not show EEG power difference between the post-test and pre-test, even though the general group power elevation was significant (see Fig. 2 & 3). A potential explanation for this phenomenon was that the MBSR intervention was not effective in some of the participants, and therefore no neurophysiological change was introduced. The same phenomenon was also found in our previous mindfulness study on the bereaved individuals (Feng-Ying Huang et al., 2021) that some of them had their mindfulness levels decreased after the 8-week mindfulness intervention. The phenomenon suggested that mindfulness training did not guarantee success for everyone. Further studies are required to figure out the reason underneath to improve the efficacy of the mindfulness training courses.

### 4.3 Situational practice effect after the 8-week training

Another focus of this study was to examine whether different mindfulness practices contribute to participants’ neuro-electrical responses differently. Compared with ***resting 1***, the MBSR-trained participants showed significantly decreased EEG activities across delta, low-beta, and parts of high-beta and gamma bands at both Fz and Pz when practicing ***body-scan***. During the body-scan practice, practitioners were instructed to be aware of their introspective body sensations. As suggested in a previous study (Mirams, Poliakoff, Brown, & Lloyd, 2013), body-scan exercise can improve people’s somatic signal detection clarity. The EEG changes associated with the ***body-scan*** task found in this study reflected how the body-scan exercise is specific and concrete for practitioners. By contrast, no significant power difference was found when practicing mindful ***breathing*** (Fig. 5), suggesting that ***body-scan*** leads to more significant EEG change among the participants than does ***breathing***.

Notably, our preliminary data collected from five mindfulness practitioners with more than two years of MBSR experience (i.e., the expert group; see Supplementary analysis, Fig. 6) further revealed that experts’ EEG power was consistently lower than that of the MBSR practitioners with 8 weeks of experience, irrespective of whether they were resting, practicing mindful breathing, or body-scan practice. In general, the high-frequency bands power of the MBSR practitioners elevated and the low-frequency bands power decreased after the 8-week MBSR training, but all EEG power decreased along with the expertise of mindfulness. Although the expert group had few participants, a general drop in EEG power across low-, middle-, and high-frequency bands could be a neuro-electrical feature of long-term MBSR practice.

**Figure 6.**
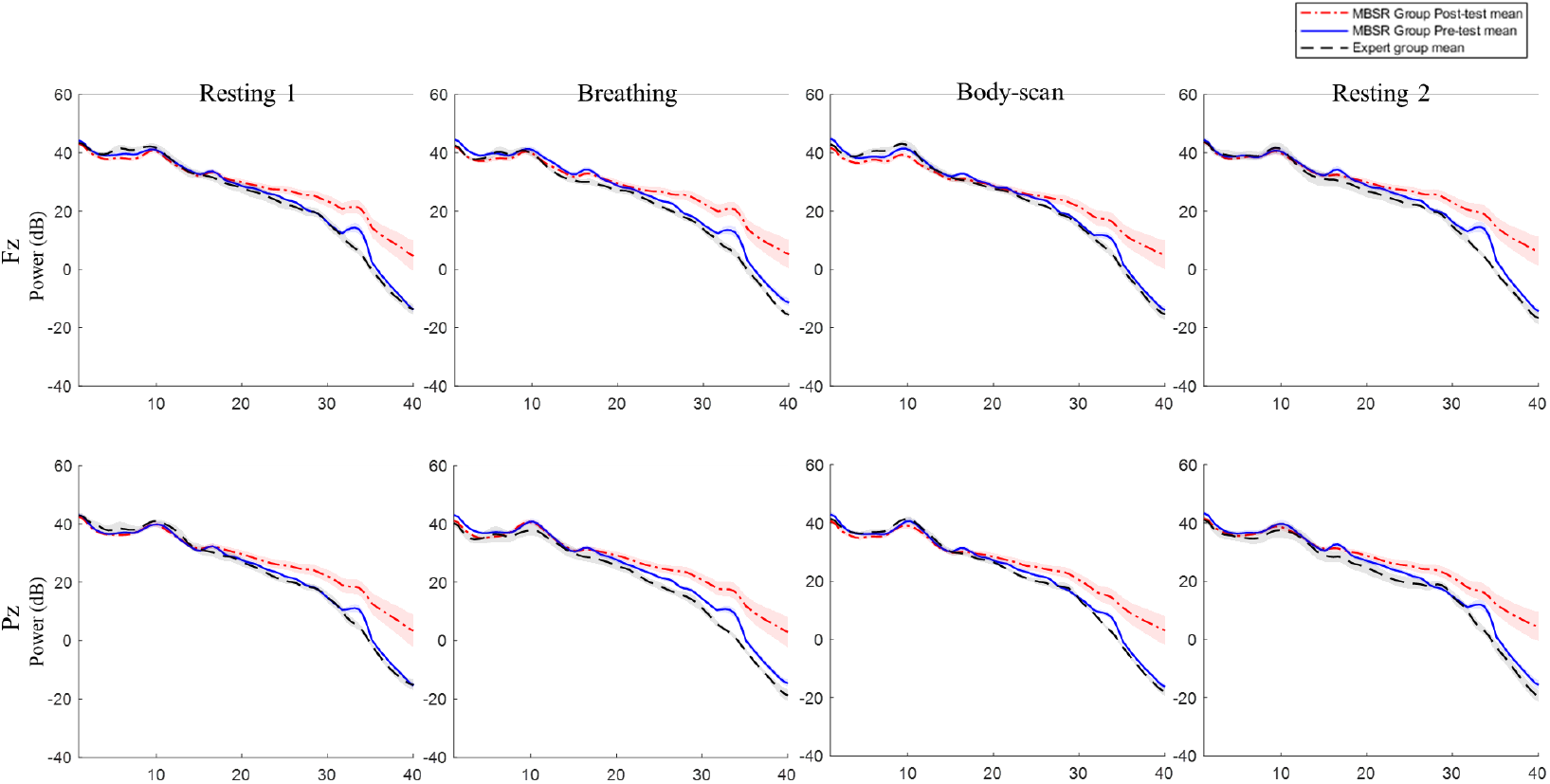
MBSR group and expert group EEG power comparison. MBSR group *n* = 17, expert group *n* = 5 Shading represents standard error of mean. No significant difference was found between expert group and the MBSR group.

### 4.4 Correlations between the change of EEG and behavioral indexes

The differences in the theta, alpha, and low-beta band powers between the post-test and pretest were correlated with the change in the FFMQ or DERS scores (as shown in Table 2). In our major findings concerning behavior and EEG biomarkers, we found significant correlations between behavioral indexes and theta, alpha, and low-beta power. These results were consistent with Ahani and colleagues’ (2014) finding that improvement in mindfulness levels was characterized by increases in the theta, alpha, and beta activities. Another study on emotional regulation also highlighted that frontal bilateral alpha activity and the parietal delta/beta ratio predicted people’s spontaneous emotion regulation level (Tortella-Feliu et al., 2014). This leads to the following question: Why was a major EEG power difference observed in high-and low-frequency bands in our data despite the clear correlation between EEG power and behavioral indexes observed in the theta, alpha, and beta bands? A possible reason could be that the major differences in neurophysiological changes after long-term mindfulness practice can be observed through delta, high-beta, and gamma spectral power and that the biomarker of mindfulness is found in alpha. A previous study on experienced Zen meditators showed that the hours of meditation practice and weekly frequency were negatively correlated with alpha power but not gamma power (Pasquini et al., 2015). Our results further suggest that mindfulness and its positive outcomes involve a complex mechanism. The EEG signals represent the mixtures of these contributing factors. Future EEG biomarker studies on mindfulness may also include the factor of cognitive ability to clarify such a complex mechanism.

### 4.5 Implication

This study demonstrated that EEG is a promising tool for probing the neurophysiological correlates of mindfulness. The evidence reported here on the long-term training effect and situational practice effect of MBSR on EEG may facilitate real-life MBSR training. For example, EEG spectral activities allow for a practical estimation of whether mindfulness-induced neural activities are activated in a practitioner. By leveraging an EEG system employing the spectral analysis, practitioners could receive neurofeedback to verify whether they are meditating correctly. Further studies on the application of EEG for facilitating mindfulness exercises will be valuable.

## Acknowledgments

This study was reviewed and approved by the Taipei Medical University Joint Institute Review Board (TMU-JIRB, project number: N201905049). This work was supported by the Center for Intelligent Drug Systems and Smart Bio-devices (IDS^2^B) from The Featured Areas Research Center Program within the framework of the Higher Education Sprout Project by the Ministry of Education (MOE), and by the Ministry of Science and Technology of Taiwan (project numbers: MOST 108-2321-B-038-005-MY2 and MOST 109-2636-E-007-022). No funding source had involved in any of the research procedures.

## CRediT authorship contribution statement

**Hei-Yin Hydra Ng**: Conceptualization, data curation, formal analysis, investigation, visualization, writing - original draft. **Changwei W. Wu**: Funding acquisition, conceptualization, project administration, supervision, writing - review & editing. **Feng-Ying Huang**: Supervision, methodology, resources - MBSR instruction. **Yu-Ting Cheng**: Data curation. **Shiao-Fei Guu**: Data curation. **Chih-Mao Huang**: Supervision, methodology. **Chia-Fen Hsu**: Supervision, methodology. **Yi-Ping Chao**: Supervision, methodology. **Tzyy-Ping Jung:** Supervision, conceptualization, methodology, writing - review & editing. **Chun-Hsiang Chuang**: Supervision, conceptualization, methodology, formal analysis, writing - review & editing.

## Appendix A

### Supplementary analysis

#### Alpha asymmetry analysis

The results of alpha asymmetry posttest pretest *t* test analysis are shown in Figure 7. The analysis method was accord with the suggestion of Isbel et al. (2019). In MBSR group, although there were trends that alpha asymmetry scores increased in ***resting 1***, the ***breathing*** and the ***body-scan*** after MBSR intervention, the differences between posttest and pretest were insignificant (*ps* > .05), neither on Fp1/2 nor F3/4. In the waitlist control group, there were no significant difference found between posttest and pretest in ***resting 1, breathing*** and ***body-scan*** (*ps* > .05), neither on Fp1/2 nor F3/4.

**Figure 7.**
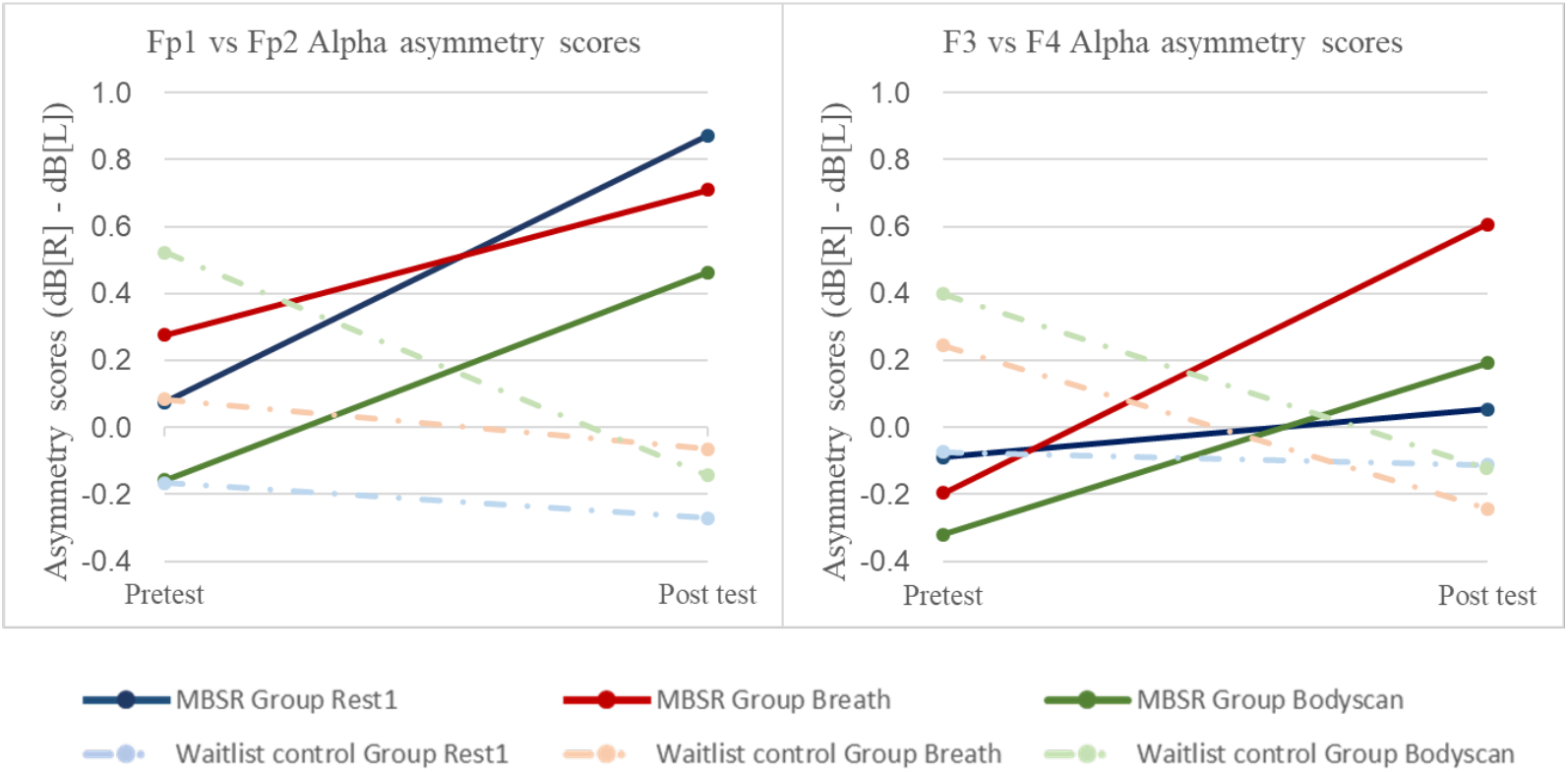
Alpha asymmetry t-test analysis on resting 1, breathing and body-scan. MBSR group *n* = 17, waitlist control group *n* = 14. No significant alpha asymmetry difference was found between posttest and pretest.

#### Between-task EEG band power differences and behavioral indexes

Table 3 shown the correlation results between the between-task EEG band power differences and behavioral indexes in the MBSR group post-test data. The results showed that the post-test MBSR group EEG spectral power difference between ***breathing*** and ***resting 1*** had a negative correlation to mindfulness level (FFMQ) and a positive correlation to difficulty of emotional regulation (DERS). No significant correlation was found between any other combination and behavioral indexes.

**Table 3.** Correlation between posttest between-task EEG band power differences and behavioral indexes in the MBSR group

| Fz | Breathing - Resting1 |  |  |  | Body-scan - Resting1 |  |  |  |
| --- | --- | --- | --- | --- | --- | --- | --- | --- |
|  | FFMQ |  | DERS |  | FFMQ |  | DERS |  |
|  | r | p | r | p | r | p | r | p |
| Delta | -0.08 | 0.747 | 0.03 | 0.895 | -0.35 | 0.166 | 0.33 | 0.199 |
| Theta | 0.04 | 0.878 | -0.25 | 0.324 | -0.23 | 0.371 | 0.12 | 0.636 |
| Alpha | 0.10 | 0.708 | -0.22 | 0.387 | -0.11 | 0.677 | 0.04 | 0.880 |
| LowBeta | -0.15 | 0.559 | -0.06 | 0.832 | -0.38 | 0.135 | 0.26 | 0.319 |
| HighBeta | 0.11 | 0.684 | -0.18 | 0.481 | 0.11 | 0.680 | -0.16 | 0.535 |
| Gamma | **-0.57\*** | 0.018 | **0.69\*** | 0.002 | -0.01 | 0.956 | <0.01 | 0.997 |
| Pz | Breathing - Resting1 |  |  |  | Body-scan - Resting1 |  |  |  |
|  | FFMQ |  | DERS |  | FFMQ |  | DERS |  |
|  | r | p | r | p | r | p | r | p |
| Delta | -0.30 | 0.245 | 0.16 | 0.549 | 0.12 | 0.658 | 0.12 | 0.650 |
| Theta | -0.19 | 0.457 | -0.05 | 0.844 | 0.11 | 0.679 | 0.02 | 0.952 |
| Alpha | -0.02 | 0.936 | -0.09 | 0.742 | 0.41 | 0.099 | -0.22 | 0.400 |
| LowBeta | -0.20 | 0.443 | 0.09 | 0.726 | 0.18 | 0.485 | 0.06 | 0.827 |
| HighBeta | -0.41 | 0.104 | 0.31 | 0.226 | 0.19 | 0.470 | <0.01 | 0.999 |
| Gamma | -0.40 | 0.111 | 0.46 | 0.061 | -0.10 | 0.713 | 0.29 | 0.265 |

*Note: \* $p < .05$*

